# Melanocyte differentiation and epidermal pigmentation is regulated by polarity proteins

**DOI:** 10.1101/2020.04.20.051722

**Authors:** Sina K. Knapp, Sandra Iden

**Affiliations:** Cologne Excellence Cluster on Cellular Stress Responses in Aging-Associated Diseases (CECAD), University of Cologne, Germany; Center for Molecular Medicine Cologne (CMMC), University of Cologne, Germany; Cell and Developmental Biology, Saarland University, Faculty of Medicine, Homburg/Saar, Germany

**Keywords:** Cell polarity, Par3, aPKC, melanocyte, differentiation, pigmentation, melanin, MITF, skin, α-MSH

## Abstract

Pigmentation serves various purposes such as protection, camouflage, or attraction. In the skin epidermis, melanocytes react to certain environmental signals with melanin production and release, thereby ensuring photo-protection. For this, melanocytes acquire a highly polarized and dendritic architecture that facilitates interactions with surrounding keratinocytes and melanin transfer. How the morphology and function of these neural crest-derived cells is regulated remains poorly understood. Here, using mouse genetics and primary cell cultures, we show that conserved proteins of the mammalian Par3-aPKC polarity complex are required for epidermal pigmentation. Melanocyte-specific deletion of Par3 in mice caused skin hypopigmentation, reduced expression of components of the melanin synthesis pathway, and altered dendritic morphology. Mechanistically, Par3 was necessary downstream of α-melanocyte stimulating hormone (α-MSH) to elicit melanin production. Strikingly, pharmacologic activation of MITF using a salt-inducible kinase inhibitor was sufficient to restore melanocyte differentiation and skin pigmentation in the absence of Par3. This data reveals a central role of polarity proteins in transmitting external pigment-inducing signals through the α-MSH/Mc1R/MITF ‘tanning pathway’, exposing unexpected links between polarity signaling and melanogenesis with new insights for pigment cell biology.

## Introduction

The mammalian skin is in constant equilibrium of regeneration and differentiation while responding to environmental cues, like pathogen attacks and UV radiation. Melanocytes ensure photo-protection of the skin by producing the pigment melanin and transferring it to neighbouring cells [1]. The interaction of melanocytes with surrounding keratinocytes, the predominant cell type in the epidermis, is vital to induce melanocyte differentiation and to trigger melanogenesis and tanning [2,3]. Altered melanocyte functions can cause hypo- and hyperpigmentation disorders [4] as well as malignant melanoma, a highly lethal skin cancer [1] Similar to cells of the peripheral nervous system, melanocytes originate from neural crest cells. Following gradual lineage specification, melanocyte precursors, so-called melanoblasts, migrate through the developing embryo along distinct yet incompletely understood paths [5]. In mice, upon crossing of the dermal-epidermal junction, melanoblasts home to the growing hair follicle or, in non-hairy skin, to the basal layer of the epidermis [4,7]. Follicular and interfollicular melanocytes coexist in the back skin epidermis during the first postnatal days. With onset of the postnatal hair follicle cycle [9], or following diverse stress signals [11,13,15], quiescent melanocyte stem cells (McSCs) are activated to provide mature, pigment-producing melanocytes. Ultimately, differentiated melanocytes assume intriguingly branched and polarized cell shapes, and possess a unique vesicular machinery for the polarized peripheral transport and concomitant maturation of melanosomes [6]. The mechanisms underlying the spatiotemporal control of melanocyte shape and plasticity, however, remain poorly understood.

Melanin production is tightly coupled to the melanocyte differentiation program. Onset of differentiation and pigmentation is marked by the activation of various pathways leading mostly to the induction of the *microphthalinia-associateA* transcription factor MITF [8]. MITF in turn drives the transcription of melanin synthesis related genes such as *Tyip1*, *Tyrp2*, *Tyr* [10] and other genes important for differentiation and survival like *cKit* [12,14,16]. Next to Notch, Wnt, endothelin and TGFß signaling [17–22], two main pathways have been reported to control the induction of MITF: the α-melanocyte stimulating hormone (α-MSH)/ melanocortin-1 receptor (Mc1R) signaling pathway, and the Stem Cell Factor (SCF)/cKit receptor tyrosine kinase pathway. As an acute physiological response to UV radiation, keratinocytes produce and secrete α-MSH, which (similar to adrenocorticotropic hormone, ACTH) can bind to Mc1R, initiating an intracellular, cAMP-mediated signaling cascade in melanocytes. Activated protein kinase A (PKA) in turn phosphorylates the cAMP-responsive-element-binding protein (CREB) [23], resulting in transcriptional activation of MITF, thereby driving skin pigmentation [8,24,25].

We and others previously demonstrated that in keratinocytes the apical polarity proteins Par3 and aPKCλ regulate cell-cell adhesion [26], stem cell fate, spindle orientation and differentiation in the epidermis [27–30]. In the adult skin epithelium, Par3 and aPKCλ promote balanced fate decisions and protect against premature epidermal differentiation, though likely through independent mechanisms [27,29,31,32]. While above work unravelled an emerging epithelial-intrinsic role of polarity proteins, we recently also uncovered extrinsic functions of polarity proteins in the control of neighbouring melanocytes: Disturbed polarity signalling in the surrounding skin epithelium resulted in aberrant keratinocyte-melanocyte adhesions, promoting melanocyte transformation and malignant progression in a melanoma mouse model [33]. Whether and how ‘apical’ polarity networks also act cell-autonomously within non-epithelial skin resident cell types is currently not known. Prompted by previous reports of polarity signalling in different neuronal processes, such as axon specification [34,35], dendritic branching [36–39], spine morphogenesis [40,41], and polarized vesicular transport [42], we here set out to investigate the significance of intrinsic polarity proteins for melanocyte function and mammalian skin pigmentation. Using different *in vivo*, *ex vivo* and *in vitro* models, we decipher a previously unrecognized causal relationship between polarity networks and melanin synthesis, with Par3 acting downstream of α-MSH to foster MITF-dependent melanogenesis.

## Results & Discussion

### Loss of Par3 and aPKC causes hypopigmentation

To explore the role of polarity signalling in melanogenesis, we first performed acute RNAi-based Par3 and aPKCλ+ζ loss-of-function in primary melanocytes isolated from P2 mice. Interestingly, knock-down of either Par3 or aPKCλ+ζ led to hypopigmented melanocytes that exhibited significantly reduced melanin levels compared to control melanocytes (Fig. 1A-C), indicating that different apical polarity proteins regulate melanocyte pigmentation. To decipher how loss of polarity signaling interferes with melanocyte function *in vivo*, we subsequently inactivated Par3 in the melanocyte lineage employing tyrosinase promoter-driven expression of Cre recombinase (Tyr-Cre [43]) combined with a conditional *Par3*^fl/fl^ allele [44,45] (Fig. S1A). Strong reduction of Par3 mRNA and protein levels was confirmed in primary melanocytes isolated from *TyrCre;Par3*^fl/fl^ (*Par3*^MCKO^) mice (Fig. S1A-C). Previous reports of melanocyte-restricted inactivation of cytoskeletal regulators like Rac1 and Cdc42 showed pigmentation defects in the coat colour (i.e. ventral patches of hypopigmented hair) due to a failure of melanoblasts to fully populate the skin in developing embryos [46,47]. Notably, at birth, *Par3*^MCKO^ mice were phenotypically indistinguishable from control littermates (Fig. S1D) and displayed a normal hair coat up to 12 months of age (Fig. S1E). This suggested that migration of melanoblast and hair coloration did not require Par3. Interestingly, however, *Par3*^MCKO^ mice showed hypopigmentation of the anterior tail skin as assessed by macroscopic analysis (Fig. 1E) and quantification of melanin content in these tissues (Fig. 1F, G). In murine tail skin-contrary to back skin-melanocytes reside not only in the hair follicles but also in distinct domains (“scales”) within the interfollicular epidermis (IFE) [48], closely resembling human skin. Melanocytes are also present in deeper dermal layers, posing the question of which melanocyte pool was responsible for the hypopigmentation in *Par3*^MCKO^ mice. Importantly, whereas melanin levels in the dermis of *Par3*^MCKO^ mice were unaltered, we found specifically the epidermal pigmentation to be reduced upon Par3 deletion (Fig. 1G). These data thus indicate that the mammalian polarity protein Par3 governs the pigmentation capability of melanocytes residing in the IFE.

**Figure 1.**
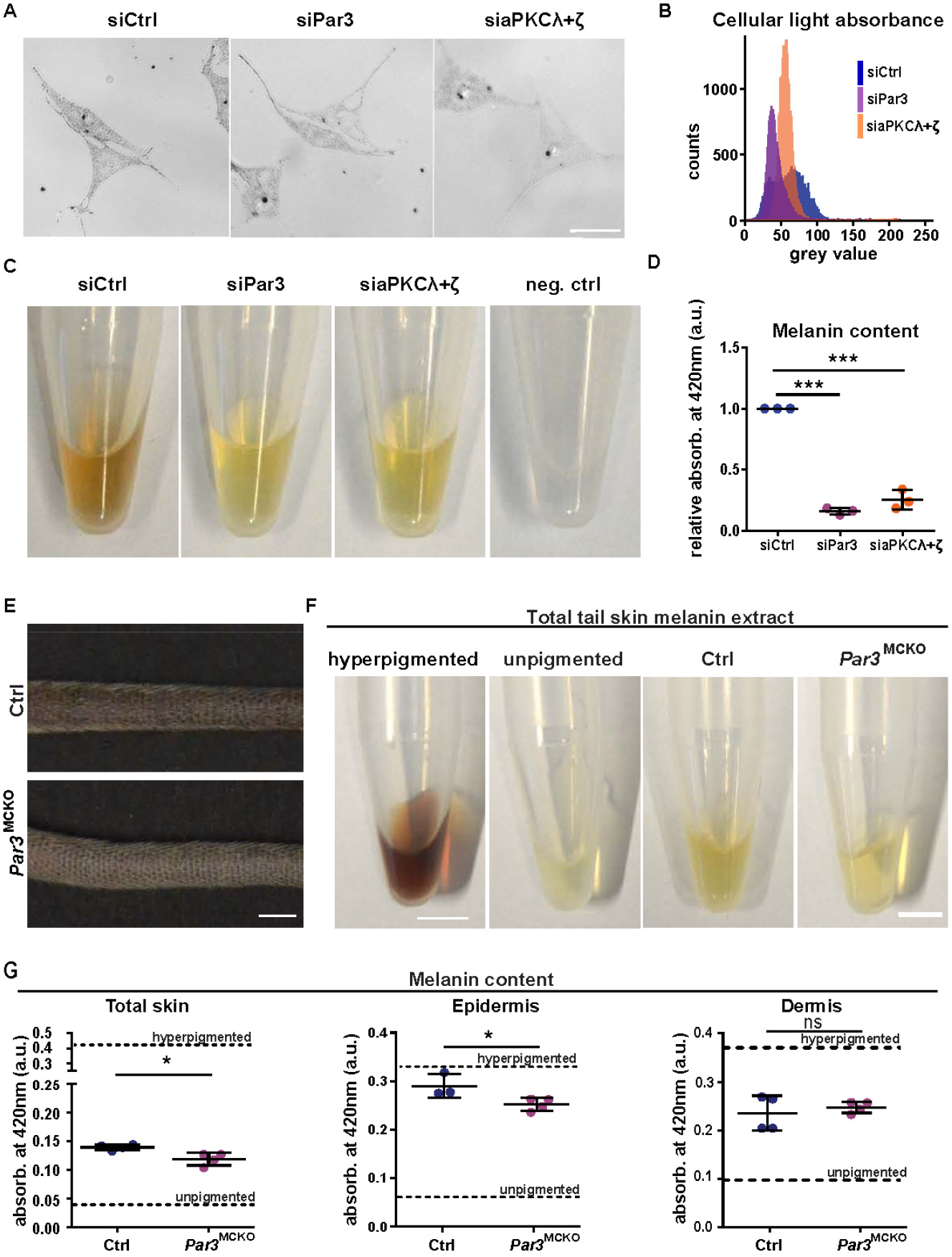
Melanocyte-specific *Par3* deletion results in tail skin hypopigmentation. A Transmitted light images of melanocytes isolated from P2 control mice transfected with nontargeting siRNA or siRNA targeting Par3 or aPKCΛ+ζ, scale bar=25μm. B Representative cellular light absorbance of a single primary melanocyte transfected with non-targeting siRNA or siRNA targeting Par3 or aPKC λ+ζ C Melanin content of primary melanocyte lysates isolated from P2 control mice transfected with non-targeting siRNA or siRNA targeting Par3 or aPKC λ+ζ CHO cells served as negative control for melanin detection, scale bar=O.5cm. D Melanin content assay. Quantification of spectrophotometry at 42Onm of siRNA treated primary melanocytes from P2 control mice, n=3, one-way ANOVA, mean±SD, ***: p<0.0001. Abbreviations: Ctrl, control; a.u., arbitrary units; P, postnatal day. E Representative image of anterior of tail from 2 months old *Par3*^MCKO^ mice and control sibling, scale bar=0.5cm. F Melanin content in tail skin lysates from 2 months old *Par3*^MCKO^ mice and littermate controls, scale bar=0.5cm. Tails of hyperpigmented mice (*Par3*^fl/fl^;HGF^tg/wt^; CDK4≡ mouse line on C57BL/6 background) and non-pigmented mice (*Par3*^fl/fl^ on FVB/N background) served as positive and negative controls, respectively. G Melanin content assay. Quantification of spectrophotometry at 420nm of total tail skin, the epidermis and the dermis of the tail skin from 2 months old *Par3*^MCKO^ and control siblings, unpaired Student’s t-tests, mean±SD, total skin: n=4, *: p=0.0136, epidermis: Ctrl: n=3, *Par3*^MCKO^: n=4, *: p=0.0426, dermis: n=4, ns: p=0.5451. Abbreviations used: Ctrl, control. See also Fig. S1.

### Par3 inactivation leads to reduced differentiation of IFE melanocytes *in vivo*

Hypopigmentation can be a result of various defects, ranging from melanocyte loss to failure to induce melanin synthesis or impaired transfer of melanosomes to the surrounding cells. In search of potential causes for the hypopigmentation in *Par3*^MCKO^ mice we therefore investigated the abundance of melanocytes *in vivo*. Notably, the number of melanocytes in scales and hair follicles was comparable to control mice (Fig. S2A), rendering it unlikely that the hypopigmentation of *Par3*^MCKO^ mice was merely a consequence of overall melanocyte loss.

In the course of differentiation, melanocytes drastically change their morphology from a fibroblast-like shape of McSCs to a pronounced dendritic appearance. Analyses of *Par3*^MCKO^ melanocytes in the tail epidermis did not reveal differences in dendricity though; with on average three major dendrites per cell in both control and *Par3*-deficient melanocytes (Fig. S2B). Notably, however, both the mean cell area and the main axis length of melanocytes in tail epidermis scales were decreased in *Par3*^MCKO^ mice (Fig. 2A, B), indicating reduced dendritic outgrowth and hence potentially reduced differentiation of *Par3*^MCKO^ scale melanocytes compared to controls. Thus far no reliable tools for endogenous markers have been established that could serve to label the stem cell pool. Instead, McSCs are typically characterized by their shape and lack of melanin, in addition to low expression of *MITF* and *cKit* [12]. Interestingly, next to their altered morphology, *Par3*^MCKO^ melanocytes in the tail scale exhibited a reduced immunoreactivity for cKit (Fig. 2C, D), supporting the concept of *Par3*^MCKO^ melanocytes residing in a less differentiated state compared to control melanocytes. These results were also interesting in light of recent data by us and others obtained in keratinocytes, where Par3 complex proteins serve to maintain stemness features of epithelial cells [27,29–31]. Our findings presented here instead uncover that in neural-crest derived melanocytes ‘apical’ polarity proteins promote differentiation, rather than stemness. Together, this highlights cell-type specific roles for these conserved polarity regulators within the complex skin tissue.

**Figure 2.**
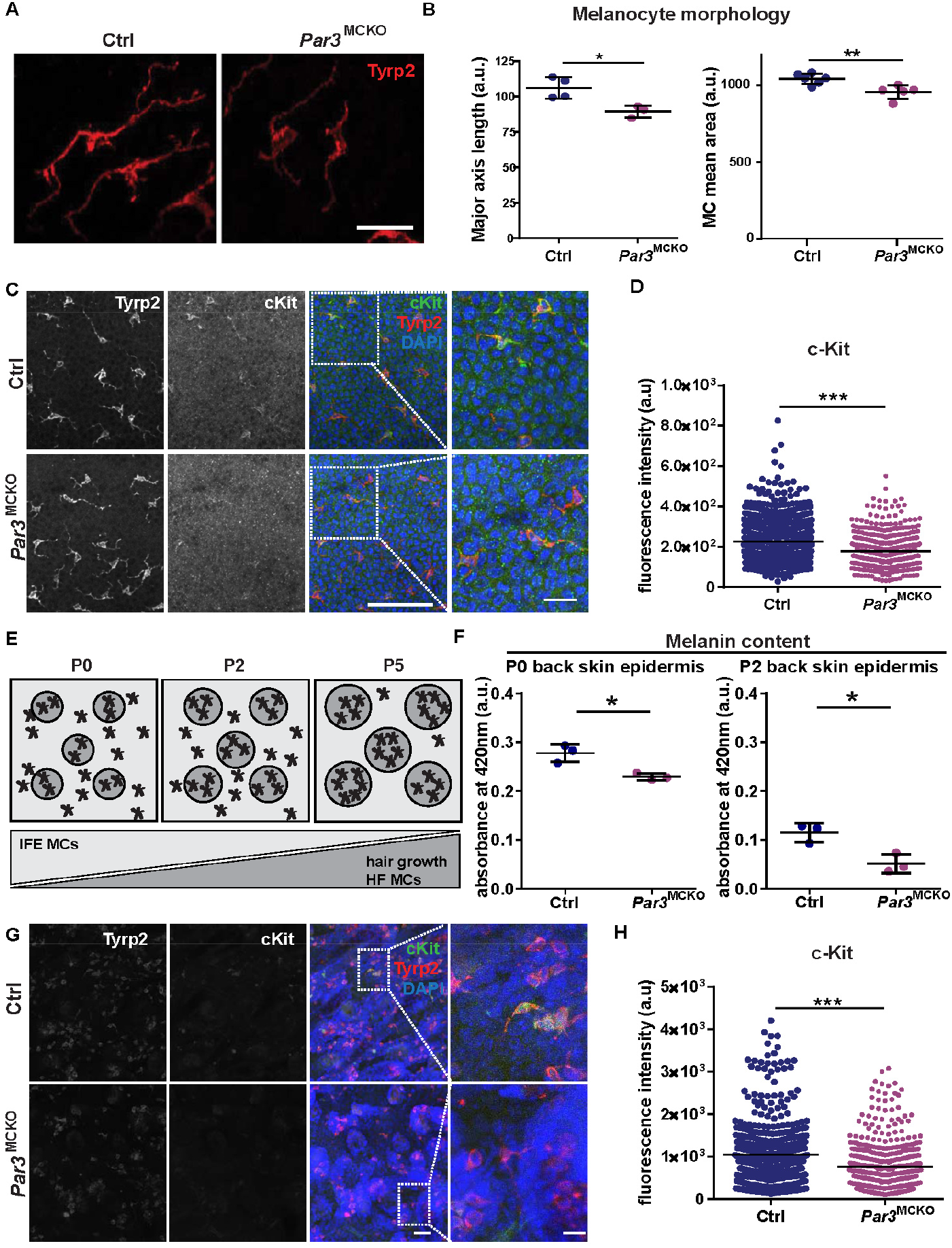
Altered differentiation of *Par3*deficient IFE melanocytes *in vivo*. A Micrograph of epidermal tail scale IFE from 2 months old *Par3*^MCKO^ and control mice stained for Tyrosinase-related protein 2 (Tyrp2). Scale bar=25μm. B Quantification of major axis length and mean area per cell of melanocytes of tail scale IFE from 2 months old *Par3*^MCKO^ and control mice, unpaired Student’s t-test, mean±SD, major axis length: Ctrl: n=4, *Par3*^MCKO^: n=3, *: p=0.0194, mean area: Ctrl: n=6, *Par3*^MCKO^: n=5, **: p=0,0057 C Micrograph of tail scale IFE melanocytes from 2 months old *Par3*^MCKO^ and control mice stained for cKit, Tyrp2 and DAPI, scale bar=30μm, magnification: scale bar=75μm, magnification: scale bar=25μm. D Quantification of cKit immunoreactivity per cell of tail scale IFE melanocytes from 2 months old *Par3*^MCKO^ and control mice, n=4, unpaired Student’s t-test, mean, ***: p<0.0001. E Scheme showing melanocyte distribution in IFE versus hair follicles age-dependently from postnatal day 0 until postnatal day 5. F Quantification of spectrophotometry at 420nm of epidermal back skin from P0 and P2, unpaired Student’s t-test, mean±SD, P0: n=3 *: p=0.0121, P2: n=3, *: p=0.0154. G Representative images from P2 back skin stained for cKit, Tyrp2 and DAPI, scale bar=75μm, magnification: scale bar=25μm. H Quantification of cKit immunoreactivity per cell, n=4, unpaired Student’s t-test, mean, ***: p<0.0001. Abbreviations: Ctrl, control; IFE, interfollicular epidermis; a.u., arbitrary units; P, postnatal day. See also Fig. S2.

### Specific requirement of Par3 for melanocytes residing in the IFE

Par3 loss selectively affected the pigmentation of adult tail IFE but not hair coloration. We therefore asked whether Par3 was also required for pigmentation of the developing epidermis in neo- and early postnatal back skin, considering its resemblance to IFE scales with respect to resident intra-epidermal melanocytes (Fig. 2E). Similar to adult tail epidermis, the pigmentation of *Par3*^MCKO^ epidermis was reduced at P0 and P2 when compared to control littermates (Fig. 2F). Similarly, the cKit immunoreactivity of back skin melanocytes in two-day old *Par3*^MCKO^ mice was lower than in controls, indicative of reduced differentiation of IFE melanocytes (Fig. 2G, H). At P5 we did not anymore detect an overt pigmentation of back skin epidermis in control and *Par3*^MCKO^ mice (Fig. S2D), in line with the developmental decline in IFE melanocytes at this stage (Fig. 2E) [49]. Moreover, no aberrant pigmentation of the dermal compartments in *Par3*^MCKO^ mice was seen at any of these stages (Fig. S2D), congruent with our observation in the dermis of adult tail skin and thus emphasizing the requirement of Par3 for melanocytes residing in close vicinity to IFE epithelial cells. The reduced cKit mean fluorescence intensity of epidermal *Par3*^MCKO^ melanocytes was further confirmed by FACS analysis of epidermis from adult tail skin and P2 back skin (Fig. S2E). We were intrigued by this distinct requirement of Par3 for the pigmentation of the IFE. Though mammalian skin epidermis and the hair follicle represent a continuous tissue, the structural and functional properties and tasks of the barrier-forming IFE and the cycling hair follicle “mini-organ” [50,51] differ significantly. For example, cells are exposed to different environmental cues dependent on their niche. UV-B radiation e.g. mainly affects cells of the IFE, whereas pathogens mainly enter the skin through hair follicles [52]. Kasper and colleagues [53] reported spatial expression signatures of the murine IFE and follicular epithelium. Of note, a recent single cell expression profiling study extended this view to distinct signatures of human IFE vs. follicular melanocytes [54], thereby confirming the previously proposed melanocyte heterogeneity [55,56]. Albeit the exact underlying mechanisms remain open, it seems likely that local parameters within the IFE, and potentially distinct keratinocyte-melanocyte interactions in that niche, contribute to the observed Par3-dependent melanogenesis.

### Suppression of melanin synthesis pathway in primary *Par3^MCKO^* melanocytes

To investigate the underlying mechanisms responsible for the hypopigmentation in epidermal tissue of *Par3*^MCKO^ mice, we isolated primary melanocytes from the back skin of P2 *Par3*^MCKO^ mice and cultured them *in vitro*. *Par3*^KO^ melanocytes contained significantly less melanin compared to control cells (Fig. 3A, B), resembling melanocytes transiently depleted of polarity proteins (Fig. 1A-C). Interestingly, *Par3*^KO^ melanocytes also showed strongly reduced RNA and/or protein levels of differentiation and pigmentation markers like Tyr, Tyrp1 and cKit (Fig. 3C-G). Moreover, nuclear immunoreactivity for the transcription factor MITF was significantly decreased in *Par3*^KO^ melanocytes (Fig. 3H, I). Notably, in contrast to epithelial cell systems in which Par3 loss has been linked to induction of apoptosis [45,57], we did not obtain evidence for increased apoptosis in *Par3*^KO^ melanocytes (Fig. S3A). Expression analysis for markers of other neuroectoderm-derived lineages such as the proneural transcription factor *Achaete-scute homolog 1* (ASCL1) [58] did not indicate transdifferentiation towards neuronal lineages following Par3 deletion (Fig. S3B). Together, these results indicate that Par3 loss in melanocytes impairs the expression of key components of the melanin synthesis pathway.

**Figure 3.**
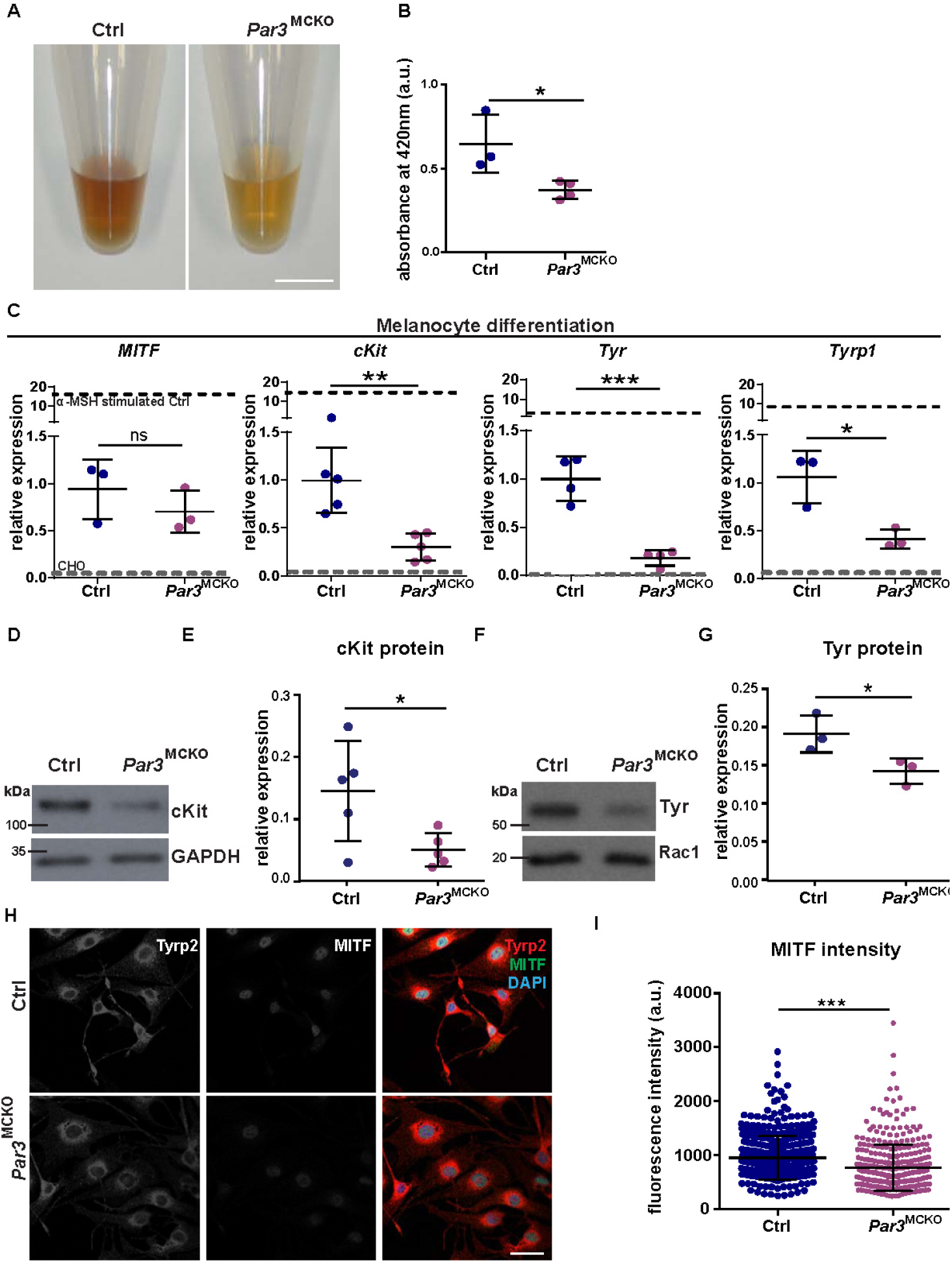
Reduced differentiation and expression of melanin synthesis genes in primary *Par3*^MCKO^ melanocytes. A Melanin content in primary melanocytes isolated from P2 *Par3*^MCKO^ and control mice, scale bar=0.5cm. B Quantification of spectrophotometry at 420nm of primary melanocyte lysates from P2 *Par3*^MCKO^ and control mice, control: n=3, *Par3*^MCKO^: n=4, unpaired Student s t-test, mean±SD, *: p=0.0280. C Quantification of RT-qPCR for *Mitf cKit*, *Tyr* and *Tyrp1* expression in primary melanocytes isolated from P2 *Par3*^MCKO^ and control mice, unpaired Student’s t-test, mean±SD, *Mitf* n=3, ns: p=0.3477; *cKit*: n=5, **: p=0.0030; *Tyr* n=4, ***: p=0.0005; *Tyrp1*: n=3, ^D^ *: p=0.0188. D Western Blot analysis of cKit expression of primary melanocytes isolated from P2 *Par3*^MCKO^ and control mice, Rac1 served as loading control. E Quantification of D), n=5, unpaired Student’s t-test, mean±SD, *: p=0.0381. F Western Blot analysis of Tyr expression of primary melanocytes isolated from P2 *Par3*^MCKO^ and control mice, GAPDH j served as loading control. G Quantification of F), n=3, unpaired Student’s t-test, mean±SD, *: p=0.0455. H Micrograph of primary melanocytes isolated of P2 *Par3*^MCKO^ and control mice immunostained for Tyrp2, MITF and DAPI, scale bar=25μm. I Quantification of MITF immunoreactivity in primary melanocytes isolated from P2 *Par3*^MCKO^ and control mice, n=3, Mann-Whitney U-test, mean, ***: p<0.0001. Abbreviations: Ctrl, control; a.u., arbitrary units; P, postnatal day. See also Fig. S3.

### Requirement of Par3 for α-MSH-induced pigmentation

We next set out to assess through which mechanisms Par3 promotes melanocyte differentiation and pigmentation. First, we asked if loss of Par3 function affected the cellular response to stimulation of the α-MSH/Mc1R pathway, a key endocrine signal inducing pigmentation. Control melanocytes, consistent with previous reports [59,60], reacted to α-MSH treatment with reinforced pigmentation (Fig. 4A, B). Interestingly, however, α-MSH was unable to increase melanin levels in *Par3*^KO^ melanocytes (Fig. 4A, B), revealing a requirement of Par3 for α-MSH-induced melanin synthesis. In contrast, we did not obtain evidence for altered TGFß or Wnt signaling in Par3-deficient melanocytes (Fig. S4A). Together, this data indicates that Par3 fosters pigmentation downstream of α-MSH.

**Figure 4.**
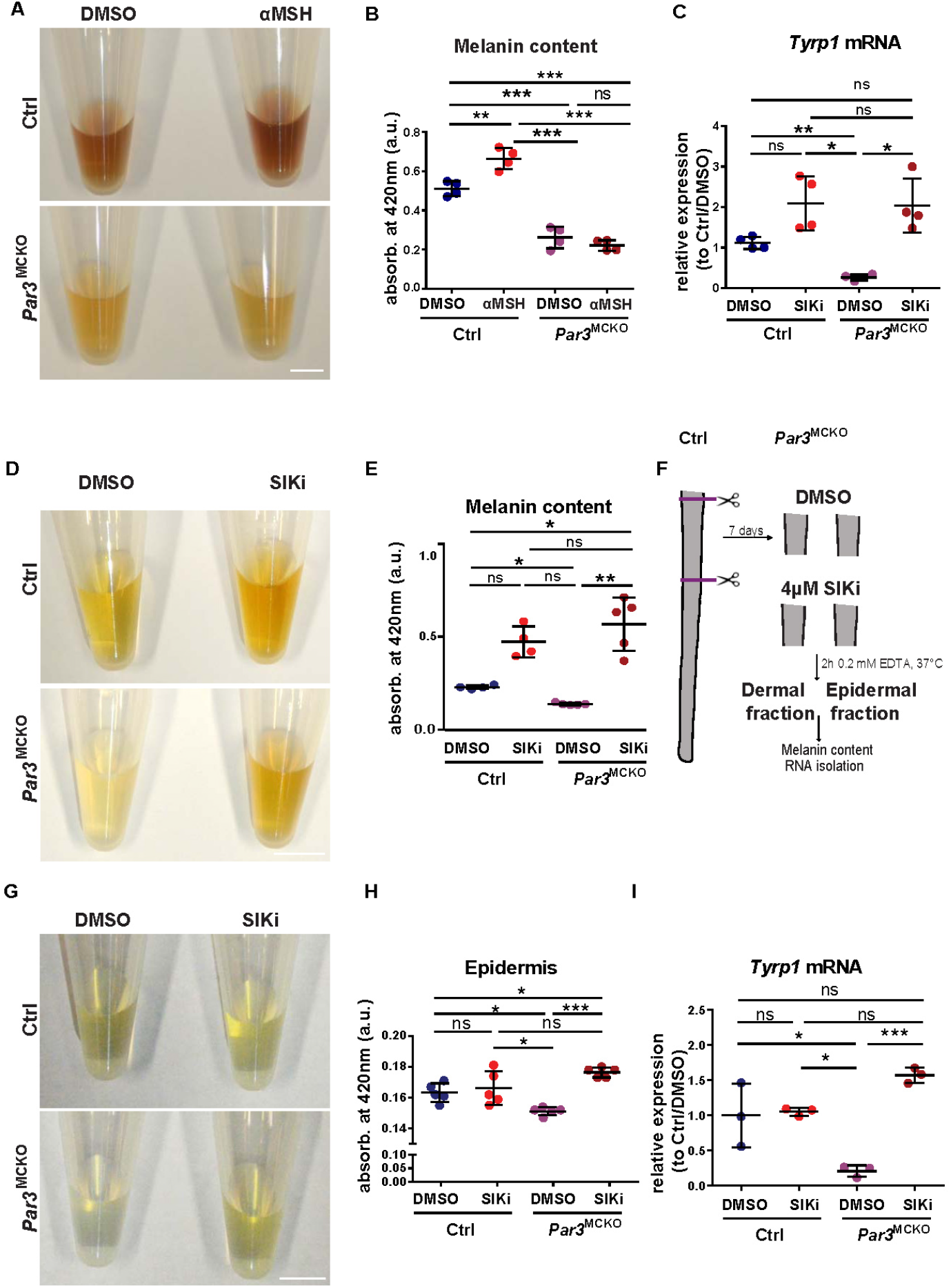
Par3 regulates MITF-induced melanocyte differentiation downstream of α-MSH. A Melanin content of lysates from primary melanocytes isolated from P2 *Par3*^MCKO^ or control mice treated with DMSO or α-MSH, scale bar=0.5cm. B Melanin content assay. Quantification of spectrophotometry at 420nm of primary melanocytes from P2 *Par3*^MCKO^ or control mice treated with DMSO or α-MSH, n=4, one-way ANOVA, mean±SD, **.: p=0.0021 (Ctrl DMSO vs Ctrl α-MSH), ***: p<0.0001 (Ctrl DMSO vs *Par3*^MCKO^ DMSO), ***: p<0.0001 (Ctrl DMSO vs *Par3*^MCKO^ α-MSH), ***: p<0.0001 (Ctrl α-MSH vs *Par3*^MCKO^ DMSO), ***: p<0.0001 (Ctrl αMSH vs *Par3*^MCKO^ α-MSH), ns: p=0.5928 (*Par3*^MCKO^ DMSO vs *Par3*^MCKO^ α-MSH). C Quantification of RT-qPCR of *Tyrp1* expression on RNA level of primary melanocytes isolated from P2 *Par3*^MCKO^ and control mice treated with DMSO or SIK inhibitor (SIKi), one-way ANOVA, n=4, mean±SD, ns: p=0.2546 (Ctrl DMSO vs Ctrl SIKi), **: p=0.0019 (Ctrl DMSO vs *Par3*^MCKO^ DMSO), ns: p=0.2499 (Ctrl DMSO vs *Par3*^MCKO^ SIKi), *: p=0.0437 (Ctrl SIKi vs *Par3*^MCKO^ DMSO), ns: p=0.9948 (Ctrl SIKi vs *Par3*^MCKO^ SIKi), *: p=0.00408 (*Par3*^MCKO^ DMSO vs *Par3*^MCKO^ SIKi). D Melanin content of primary melanocytes’ lysates isolated from P2 *Par3*^MCKO^ and control mice treated with DMSO or SIKi, scale bar=0.5cm. E Quantification of D. Spectrophotometry at 420nm of primary melanocyte lysates from P2 *Par3*^MCKO^ and control mice, n=4, mean±SD, *: p=0.0295 (Ctrl DMSO vs Ctrl SIKi), **: p=0.0030 (Ctrl DMSO vs *Par3*^MCKO^ DMSO), ns: p=0.052 (Ctrl DMSO vs *Par3*^MCKO^ SIKi), *: p=0.0128 (Ctrl SIKi vs *Par3*^MCKO^ DMSO), ns: p=0.7822 (Ctrl SIKi vs *Par3*^MCKO^ SIKi), *: p=0.0339 (*Par3*^MCKO^ DMSO vs *Par3*^MCKO^ SIKi). F Scheme explaining *ex vivo* SIKi treatment of anterior tail skin, treated either with DMSO or SIKi for 7 days, followed by separation of epidermal and dermal fraction, and melanin content assay or RNA analysis. G Melanin content of anterior tail skin epidermis’ lysates treated with DMSO or SIKi, scale bar=0.5cm. H Quantification of spectrophotometry at 420nm of anterior tail skin epidermis treated with DMSO or SIKi from *Par3*^MCKO^ and control mice, n=5, one-way ANOVA, mean±SD, ns: p=0.5893 (Ctrl DMSO vs Ctrl SIKi), *: p=0.0258 (Ctrl DMSO vs *Par3*^MCKO^ DMSO), *: p=0.0302 (Ctrl DMSO vs *Par3*^MCKO^ SIKi), ns: p=0.1118 (Ctrl SIKi vs *Par3*^MCKO^ DMSO), ns: p=0.2810 (Ctrl SIKi vs *Par3*^MCKO^ SIKi), **: p=0.0011 (*Par3*^MCKO^ DMSO vs *Par3*^MCKO^ SIKi). I Quantification of RT-qPCR for *Tyrp1* expression of anterior tail skin epidermis treated with DMSO or SIKi from *Par3*^MCKO^ and control mice, n=3, one-way ANOVA, mean±SD, ns: p=0.9933 (Ctrl DMSO vs Ctrl SIKi), *: p=0.0149 (Ctrl DMSO vs *Par3*^MCKO^ DMSO), ns: p=0.0730 (Ctrl DMSO vs *Par3*^MCKO^ SIKi), *: p=0.0106 (Ctrl SIKi vs *Par3*^MCKO^ DMSO), ns: p=0.1054 (Ctrl SIKi vs *Par3*^MCKO^ SIKi), ***: p=0.0011 (*Par3*^MCKO^ DMSO vs *Par3*^MCKO^ SIKi). Abbreviations: Ctrl, control; a.u., arbitrary units; P, postnatal day; SIKi, SIK inhibitor. See also Fig. S4.

### Par3 acts upstream of MITF signaling to promote melanin synthesis

To further delineate the role of Par3 in melanin synthesis, we stimulated primary *Par3*^KO^ melanocytes with HG 9-91-01, an inhibitor of saltinducible kinase (SIK). SIK antagonizes MITF transcription by phosphorylating CRTC, thereby preventing its nuclear translocation and as consequence the activation of CREB [61]. PKA instead is a positive regulator of MITF, counteracting SIK function. SIK inhibition by HG 9-91-01 has recently been reported as a UV-independent approach to induce MITF-driven skin pigmentation [62]. Intriguingly, SIK inhibitor treatment of primary *Par3*^KO^ melanocytes was sufficient to restore *Tyrp1* expression (Fig. 4C) and melanin content to the level of control melanocytes (Fig. 4D, E). Strikingly, next to this reinstalled pigmentation of Par3-deficient melanocytes *in vitro*, SIK inhibition also rescued the hypopigmentation of *Par3*^MCKO^ explants (Fig. 4F, G, H). Differential melanin content analysis revealed that this was due to a significant increase of melanin in the epidermal compartment (Fig. 4G, H), whereas pigmentation of the corresponding dermal explant fraction was unaltered (Fig. S4B). In line with the restored epidermal melanin levels, *Tyrp1* expression in the epidermal fraction of SIK inhibitor-treated *Par3*^MCKO^ skin was significantly elevated and comparable to that of control epidermis (Fig. 4I). These results thus suggest that Par3 promotes epidermal melanocyte pigmentation and differentiation through steering of MITF function. Considering the emerging albeit controversial roles of MITF in melanoma formation, progression and therapeutic resistance [4,69,70], our findings also raise new questions about the relevance of Par3/MITF signaling in modulating melanoma plasticity.

Collectively, this study revealed unexpected links between mammalian cell polarity proteins and the melanin synthesis pathway important to mediate skin pigmentation. By combining mouse genetics, primary cell systems, quantitative imaging and gene expression analyses, we showed that the polarity proteins Par3 and aPKC serve as crucial determinants for melanogenesis. Loss- and gain-of-function approaches delineate a role of Par3 downstream of the α-MSH/Mc1R pathway, governing the expression of key components of the melanin synthesis pathway and melanocyte differentiation. Pharmacologic rescue of the hypopigmentation in *Par3*^MCKO^ melanocytes and epidermal tissues by a SIK inhibitor further demonstrated that Par3 acts upstream of MITF, a master transcription factor in the melanocyte lineage (Fig. 5A-C). Together, this work unravels a requirement of polarity signaling for melanocyte differentiation and function, providing an important basis for future mechanistic investigation of mammalian skin pigmentation and pathomechanisms underlying pigmentation disorders.

**Figure 5.**
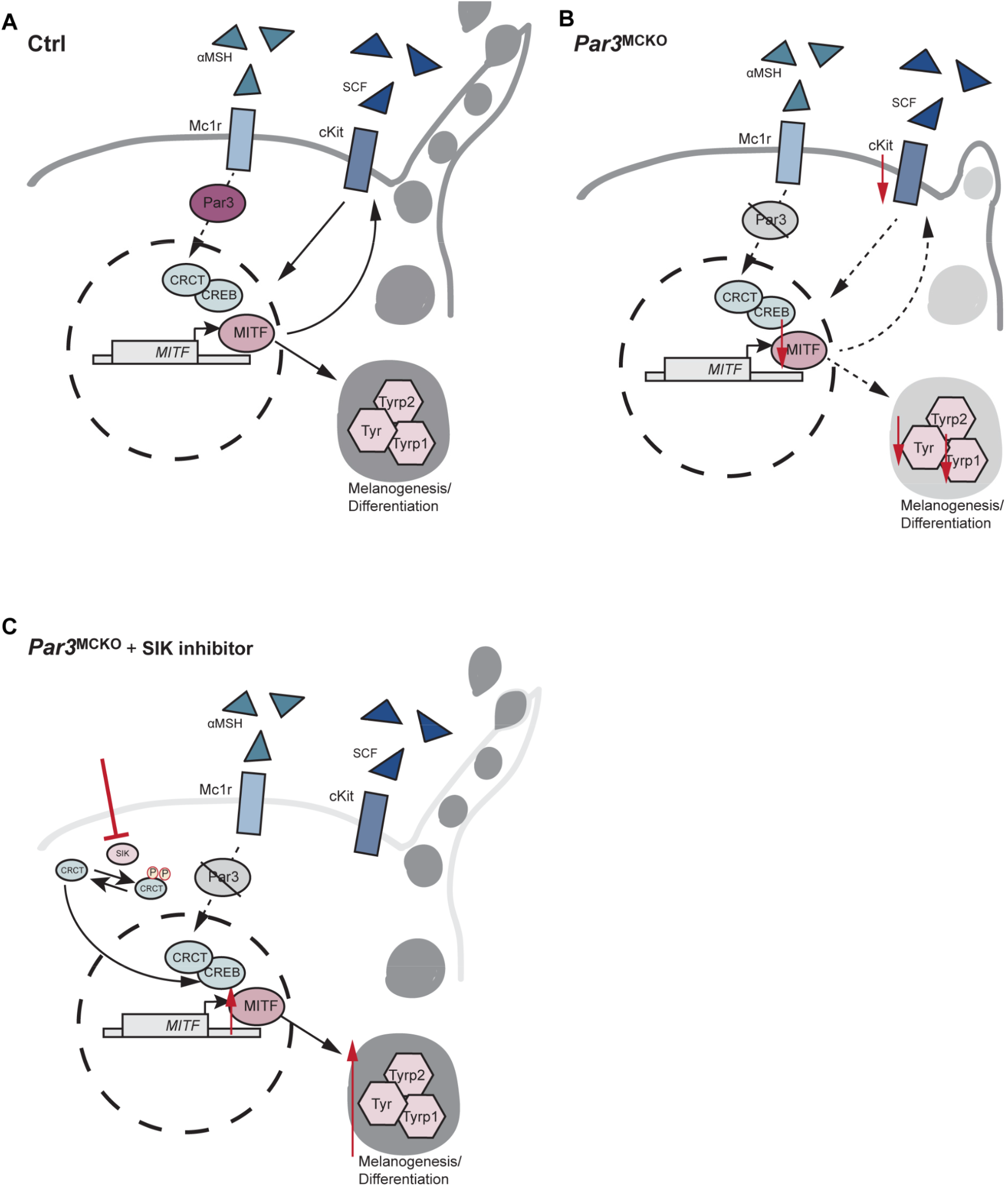
Working model. Compared to control melanocytes (A), Par3-depleted melanocytes are hypopigmented and less differentiated *in vivo* and *in vitro*, and do not induce melanogenesis following α-MSH treatment (B). Components of the melanin synthesis pathway, and resulting pigmentation, are restored upon treatment with a SIK inhibitor in the absence of Par3 (C), identifying this polarity protein as a key regulator of melanocyte homeostasis upstream of MITF.

## Experimental Procedures

### Mice

*Tyrosinase-Cre* mice [43] were crossed with *Par3* flox mice [44]. Animals were bred on C57BL/6J background and were fed and housed according to federal guidelines. All animal experiments were performed according to institutional guidelines and animal licenses by the State Office of North Rhine-Westphalia (LANUV), Germany. Specific primers for genotyping are listed in Table S3.

### Melanocyte isolation and culture

Primary mouse melanocytes were isolated from the epidermis of newborn control and *Par3*^MCKO^ mice. To separate epidermis from dermis, whole skins of postnatal day 2 (P2) mice were incubated in 5mg/ml Dispase II (Sigma-Aldrich) diluted in RPMI medium supplemented with 10% FCS, penicillin (100U/ml), streptomycin (100μg/ml, Biochrom), 100nM sodium pyruvate, and 10mM non-essential amino acids (all Gibco) at 4°C overnight (RPMI medium hereafter). Epidermis was separated from the dermis and incubated for 20 min in TrypLE (Gibco) at room temperature before dissociated cells were collected and cultured in RPMI medium containing 200nM TPA and 200pM cholera toxin (RPMI+ hereafter) (both Sigma-Aldrich). At passage 0, cultures contained melanoblasts, melanocytes and keratinocytes. After 7 days, cells were passaged and due to terminal differentiation, keratinocytes were largely excluded, resulting in melanocyte monocultures.

### Immunocytochemistry

Cells grown on glass chamber slides or permanox were fixed in 4% paraformaldehyde (PFA) or EtOH/acetone, permeabilized with 0.5% Triton-X in PBS (if PFA fixed) and blocked in 1% Triton X/10% fetal calf serum (FCS) in PBS. Samples were incubated overnight in primary antibodies diluted in blocking buffer, followed by washing and incubation in secondary antibodies for 1h at room temperature. Finally, samples were mounted in Mowiol. Antibodies used for stainings are listed in Table S1.

### Isolation and immunohistochemistry of tail epidermis whole mounts

Whole mounts of tail skin and back skin epidermis were prepared as previously described [63]. In brief, tail epidermis was separated from dermis after 4h incubation in 5mM EDTA/PBS at 37°C and fixed in 4% PFA/PBS for 1h at room temperature. Tail skin epidermis was blocked for 1h in PB buffer (0.9% NaCl and 20 mM HEPES, pH 7.2, plus 0.5% skim milk powder, 0.25% fish skin gelatin (Sigma-Aldrich), and 0.5% Triton X-100), back skin epidermis was blocked for 1h in 10% FCS plus 1% Triton X in PBS at RT. Primary antibody incubation was performed in the respective blocking buffer at room temperature overnight. After washing 3x in 0.2% Tween/PBS for 1h, whole mounts were stained with secondary antibodies at room temperature overnight. After washing, whole mounts were mounted in Mowiol. Antibodies used for stainings are listed in Table S1.

### Preparation of cell extracts, SDS-PAGE, and immunoblotting

Melanocyte cultures were lysed with crude lysis buffer (10mM EDTA, 1% SDS), protein concentrations were determined via BCA assay (Pierce, Thermo Fisher Scientific Darmstadt, Germany), and SDS-PAGE (8-12% PAA) and immunoblotting was performed according to standard procedures [26].

### Microscopy

Confocal images were acquired with a Leica SP8 and a Zeiss Meta 710 laser scanning microscope using a Plan-Apochromat 63x/1.4 NA oil, Plan-Neofluar 20x/0.8, Plan-Neofluar 40x/1.3 Oil DIC, 40x air. Epifluorescence images were obtained with a Leica DMI6000 and the following objectives: PlanApo 63x, 1.4 NA; PlanApo 20x, 0.75 NA.

### Quantification of MITF immunoreactivity

A Cell Profiler [64] pipeline was used to calculate MITF immunoreactivity in melanocyte nuclei. Briefly, DAPI staining was used as seeds for primary object detection. MITF signals were identified as objects and related to DAPI seeds. Analysis was done automatically with supervision, and data were exported to a spreadsheet.

### Quantification of cKit immunoreactivity

A Cell Profiler [64] pipeline was used to calculate cKit immunoreactivity in melanocytes. Briefly, Tyrp2 staining was used as seeds for primary object detection. cKit signals were identified as objects and related to Tyrp2 seeds. Analysis was done automatically with supervision, and data were exported to a spreadsheet.

### Quantification of protein signals in immunoblot analyses

The band intensity of non-saturated Western blot signals was determined using the GelAnalyzer software of scanned Western blots. Quantifications show the relative protein signals after normalization to loading controls and subsequent normalization to the sum of the densitometric signal of paired samples as described previously [65].

### Inhibitor treatment

To induce pigmentation, cells or tissues were treated with 4μM SIK inhibitor (MedChemExpress, #HY-15776) diluted in RPMI+ for 24h, 72h or 7 days. Medium with SIK inhibitor was exchanged every 24h, as described previously [62].

### α-MSH treatment

To induce differentiation and pigmentation, melanocytes were treated with 100nM α-Melanocyte Stimulating Hormone (α-MSH) (Calbiochem, #05-23-0751) for 72h. Medium with α-MSH was exchanged every 24h.

### SiRNA-mediated knockdown

To transiently down-regulate Par3 or aPKC, primary melanocytes were transfected using Viromer^®^ (Lipocalyx) with 100nM of ON-TARGETplus SmartPool targeting murine Par3, aPKC or a non-targeting control siRNA pool (siPar3: equimolar mixture of four Par3 targeting SmartPools: J-040036-05, J-040036-06, J-040036-07, J-040036-08, Dharmacon, siaPKC: equimolar mixture of aPKC λ and aPKCζ targeting smartpools: siaPKC λ: J-040822-05, J-040822-06, J-040822-07, J-040822-08, Dharmacon, siaPKCζ: J-040823-09, J-040823-10, J-040823-11, J-040823-12, Dharmacon, siCtr: D-001206-14-20, Dharmacon) following the manufacturer’s protocol. Cells were cultivated in RPMI+ medium for 48h prior to lysis. Knockdown efficiency was validated by Western blot analysis using Par3- and aPKC-specific antibodies, respectively.

### RNA isolation from primary melanocytes

Melanocytes were cultured in 6cm dishes and washed with PBS prior to lysis with 500μl TRIzol reagent (#15596, ThermoFisher Scientific). After homogenization, samples were incubated for 5 min at RT prior to addition of 100μl chloroform. Incubation for 3 min at RT followed centrifugation at 12.000 x g for 15 min at 4° C. The RNA containing aqueous phase was transferred to a new reaction tube and 500μl 100% isopropanol was added for RNA precipitation. After 10 min incubation at RT, samples were centrifuged at 12.000xg for 10 min at 4°C. Supernatants were discarded and remaining pelleted RNA was washed in 1ml 75% ethanol before centrifugation at 7500xg for 5 min at 4°C. After removal of supernatants RNA was air-dried for 5 min and resuspended in 15μl DEPC water.

### RT-qPCR

RNA was transcribed into cDNA using QuantiTect Reverse Transcription Kit^®^ (Qiagen). Different TaqMan gene expression assays^®^ (ThermoFisher Scientific), listed in Table S2, were used to perform quantitative real-time PCR. Gene expression changes were calculated using the comparative CT method and normalized to GAPDH before normalization to control cells or epidermal lysates.

### Flow cytometry analysis

P2 dorsal skin samples or P58 tail skin samples were obtained from transgenic mice. Incubation of the whole skin in 0,08% Trypsin in PBS for 50 min at 37° C allowed the separation of the epidermis from the dermis. Fragmented epidermis was incubated in RPMI medium for 30 min at 37°C and subsequently passed through 40μm nylon mesh (BD Falcon). Epidermal cell suspensions were washed in PBS prior to antibody incubation in 2% FCS, plus 2 mM EDTA in PBS (FACS buffer) for 1h at 4° C. After washing epidermal cell suspension in PBS, flow cytometry analysis was conducted on a LSRFortessa Flow Cytometer (BD Biosciences), followed by analysis with FACSDiVa software, version 8.0.1.

### Melanin content assay

Melanin content was measured in an adapted protocol from Ito and Wakamatsu [66]. Back skin, tail skin, back fur of mice, or primary melanocytes were lysed in 10% DMSO in NaOH for 24h at 100°C. Absorbance of the lysates was measured at 420 nm using a spectrophotometer (EnSpire Multimode Plate Reader, Perkin Elmer).

### Antibodies

Details on the antibodies used in this study are listed in Table S1.

### Statistical analyses

GraphPad Prism Software (GraphPad, version 6.0) was used to perform statistical analyses. Significance was determined by Mann-Whitney U-test, Student’s *t*-test, One-Way ANOVA Sidak’s multiple comparisons test, as indicated in the figure legends. All data sets were subjected to normality tests (D’Agostino-Pearson omnibus test, KS normality test, or Shapiro-Wilk normality test) when applicable. N-values correspond to the sample size used to derive statistics. P-values are ranged as follows: *, p≤0.05, **, p≤0.01, ***, p≤0.001 as detailed in the figure legends. The number and type of biological replicates is specified in the figure legends. For all experiments, measurements were taken from minimum three independent biological samples.

### Software

Data collection utilized the following software: Microscopy: LASX (Leica), ZEN (Zeiss), Volocity (PerkinElmer): Immunoblot: Samsung SCX3405 printer/scanner: GelDoc (Biorad): qRT-PCR: QuantStudio 12K Flex Software (Thermo). For data analysis, the following software has been used: GraphPad PRISM VI, ImageJ/Fiji [67,68], Photoshop, GelAnalyzer 2010, FACSDiVa software, version 8.z0.1, Cell Profiler [64].

## Data Availability

Correspondence and requests for materials related to this study should be sent to. All data supporting the findings of this study are available within the paper and its supplemental information, or from the corresponding author on reasonable request.

## Acknowledgements

We thank Michael Saynisch for technical assistance, and the CECAD imaging facility and the UoC animal facilities for important services. We thank all members of the Iden laboratory for stimulating discussions, and Mengnan Li and Annika Graband for critical reading of the manuscript. Furthermore, we are grateful to Kathleen J. Green and to the Niessen, Bazzi, and Wickström laboratories for helpful input and various discussions. We acknowledge Lionel Larue and Shigeo Ohno for sharing mouse lines. This project was funded by the Deutsche Forschungsgemeinschaft (DFG, German Research Foundation) (grants SPP1782-ID79/2-1, SPP1782-ID79/2-2, and Projektnummer 73111208 – SFB 829, A10). Work in the Iden laboratory was further supported by the Saarland University, Excellence Initiative of the German federal and state governments (CECAD Cologne), and Center for Molecular Medicine Cologne (CMMC).

## Author Contributions

Conceptualization, methodology, and validation: S.K., S.I.; investigation: S.K., S.I.; formal analysis: S.K., S.I.; resources: S.I.; visualization: S.K.; writing (original draft): S.K.; writing (review and editing): S.I.; supervision and funding acquisition: S.I.

