## Supplemental Information for "Melanocyte differentiation and epidermal pigmentation is regulated by polarity proteins"

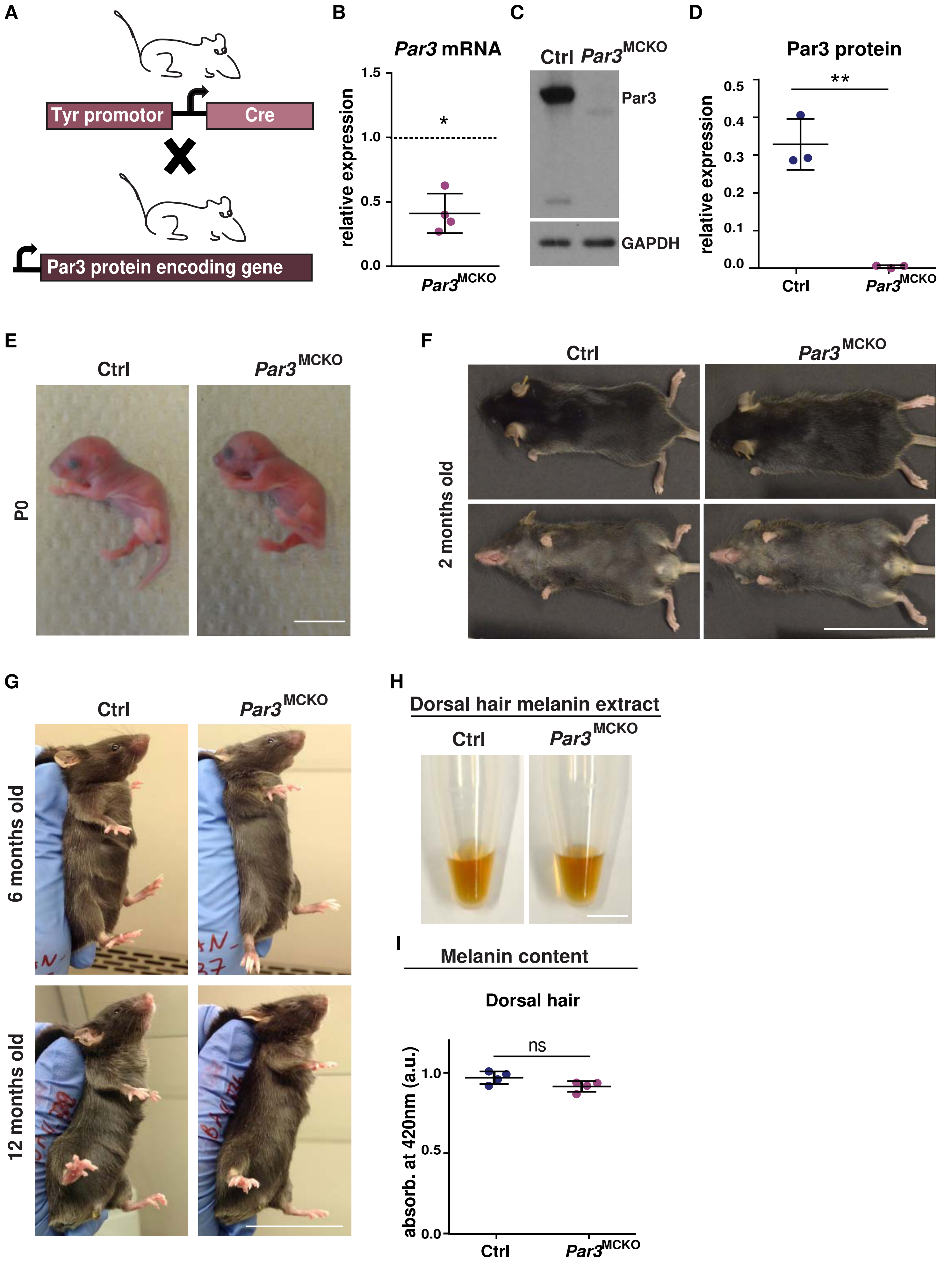

A
Tyr promotor
Cre
Par3 protein encoding gene
B
Par3 mRNA
relative expression
1.5
1.0
0.5
0.0
*
Par3MCKO
C
Ctrl
Par3MCKO
Par3
GAPDH
D
Par3 protein
relative expression
0.5
0.4
0.3
0.2
0.1
0.0
**
Ctrl
Par3MCKO
E
Ctrl
Par3MCKO
P0
F
Ctrl
Par3MCKO
2 months old
G
Ctrl
Par3MCKO
6 months old
12 months old
H
Dorsal hair melanin extract
Ctrl
Par3MCKO
I
Melanin content
Dorsal hair
absorb. at 420nm (a.u.)
1.0
0.5
0.0
ns
Ctrl
Par3MCKO

**Figure S1. Phenotypic characterization of *Par3*$^{MCKO}$ mice. Related to Figure 1. A)** Breeding scheme for the generation of melanocyte-specific knock-out for the polarity protein Par3. **B)** Quantification of RT-qPCR for *Par3* expression of isolated melanocytes from newborn *Par3*$^{MCKO}$ mice and control siblings, n=4, one-sample Student's t-test. *: p=0.0128, mean±SD. Residual Par3 expression is likely attributed to low amounts of remaining keratinocytes in melanocyte cultures. **C)** Western Blot analysis for *Par3* expression in melanocytes isolated from newborn *Par3*$^{MCKO}$ mice and control siblings. GAPDH served as loading control. **D)** Quantification of C), n=3, unpaired Student´s t-test, **: p=0.0011, mean±SD. **E)** Phenotypic characterization of P0 *Par3*$^{MCKO}$ mouse compared to control sibling, scale bar=1cm. **F)** Phenotypic characterization of 2 months old *Par3*$^{MCKO}$ mouse compared to control sibling, scale bar=5cm. **G)** Phenotypic characterization of 6 and 12 months old *Par3*$^{MCKO}$ mice compared to control siblings, scale bar=5cm, **H)** Melanin content of dorsal hair lysates from 2 months old *Par3*$^{MCKO}$ and control mice, scale bar=0.5cm. **I)** Melanin content assay. Quantification of spectrophotometry at 420nm of dorsal hair from 2 months old *Par3*$^{MCKO}$ and control mice, n=4, unpaired Student´s t-test, mean±SD, ns: p=0.0809. Abbreviations: Ctrl, control; a.u., arbitrary units; P, postnatal day.

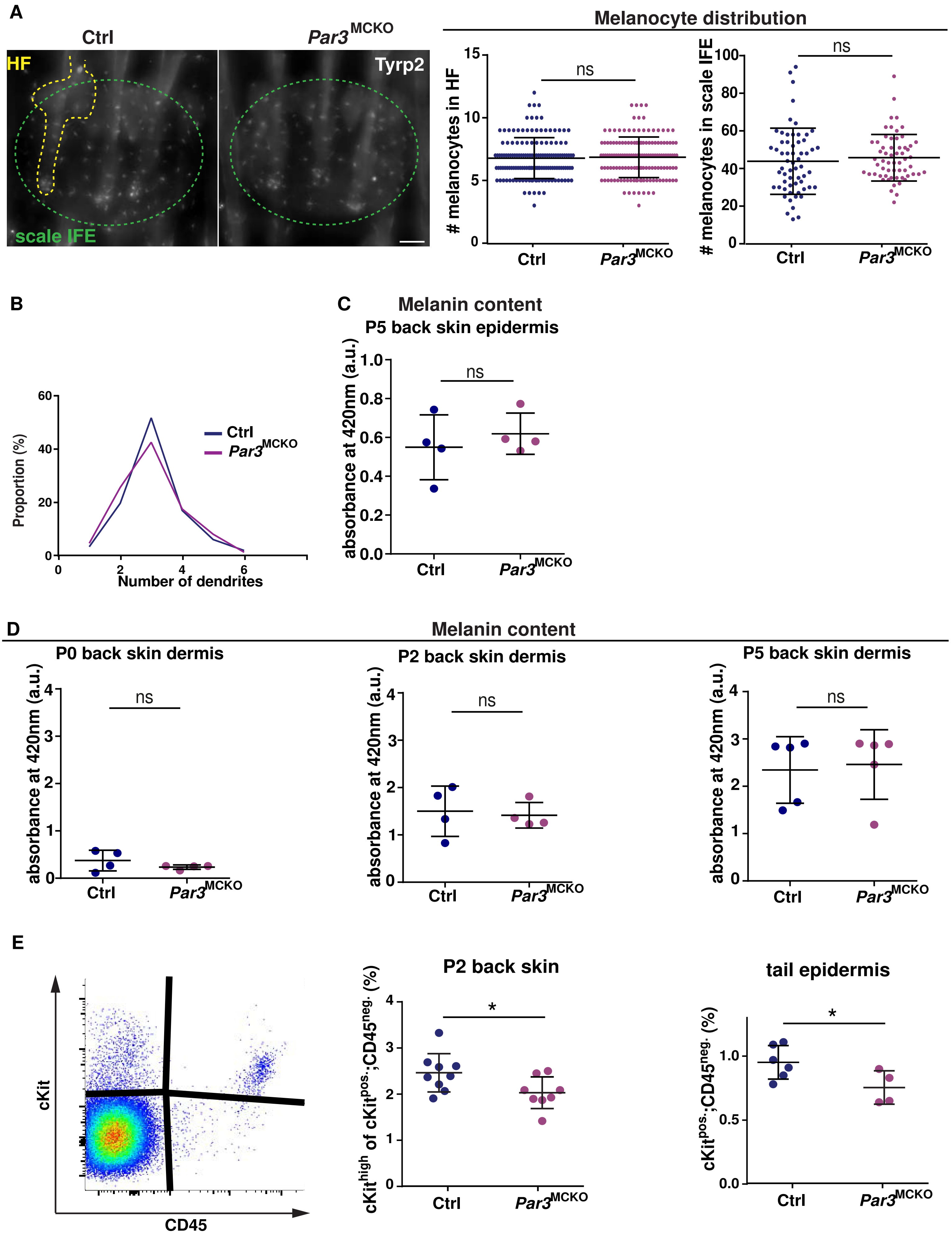
A
Ctrl
Par3MCKO
HF
Tyrp2
scale IFE
Melanocyte distribution
### melanocytes in HF
ns
Ctrl
Par3MCKO
### melanocytes in scale IFE
B
Proportion (%)
Ctrl
Par3MCKO
Number of dendrites
C
Melanin content
P5 back skin epidermis
absorbance at 420nm (a.u.)
D
Melanin content
P0 back skin dermis
P2 back skin dermis
P5 back skin dermis
E
cKit
CD45
P2 back skin
cKithigh of cKitpos.;CD45neg. (%)
*
tail epidermis
cKitpos.;CD45neg. (%)

**Figure S2. Melanocyte numbers and dermal pigmentation is not altered in *Par3*$^{MCKO}$ mice. Related to Figure 2. A)** Micrograph of melanocytes located in the scale IFE and the HF in the tail skin epidermis in 2 months old *Par3*$^{MCKO}$ and control mice, scale bar=75µm, Quantification of the respective melanocyte numbers in HF and scale IFE in 2 months old *Par3*$^{MCKO}$ and control mice, n=3, unpaired Student´s t-test, mean±SD, melanocyte number in HF: ns: p=0.7263, melanocyte number in scale IFE: ns: p=0.4910. **B)** Representation of dendrite number per cell in tail scale IFE melanocytes of 2 months old *Par3*$^{MCKO}$ and control mice. **C)** Quantification of spectrophotometry at 420nm of epidermal back skin from P5, unpaired Student´s t-test, mean±SD, P5: n=4, ns=0.5072. **D)** Melanin content assay. Quantification of spectrophotometry at 420nm of dermal back skin from *Par3*$^{MCKO}$ and control mice at the postnatal days 0, 2 and 5, unpaired Student´s t-test, mean±SD, P0: n=4, ns: p=0.2591, P2: n=4, ns: 0.7829, P5: n=5, ns: 0.8049. **E)** Left panel: Example flow cytogram of epidermal skin suspension stained with anti-CD45 (to exclude hematopoietic cells) and anti-CD117 (cKit) antibodies, right panel: Quantification of cKit MFI in cKit$^{pos}$;CD45$^{neg}$ cell population of epidermal back skin from P2 *Par3*$^{MCKO}$ and control mice and of cKit$^{pos.}$;CD45$^{neg.}$ cells from the anterior tail skin epidermis of 2 months old *Par3*$^{MCKO}$ and control mice, P2 back skin: Ctrl: n=9, *Par3*$^{MCKO}$: n=8, unpaired Student´s t-test, mean±SD, *: p=0.0350, tail epidermis: Ctrl: n=6, *Par3*$^{MCKO}$: n=4, unpaired Student´s t-test, mean±SD, *: p=0.0483. Abbreviations: Ctrl, control; a.u., arbitrary units; P, postnatal day; MFI, mean fluorescence intensity.

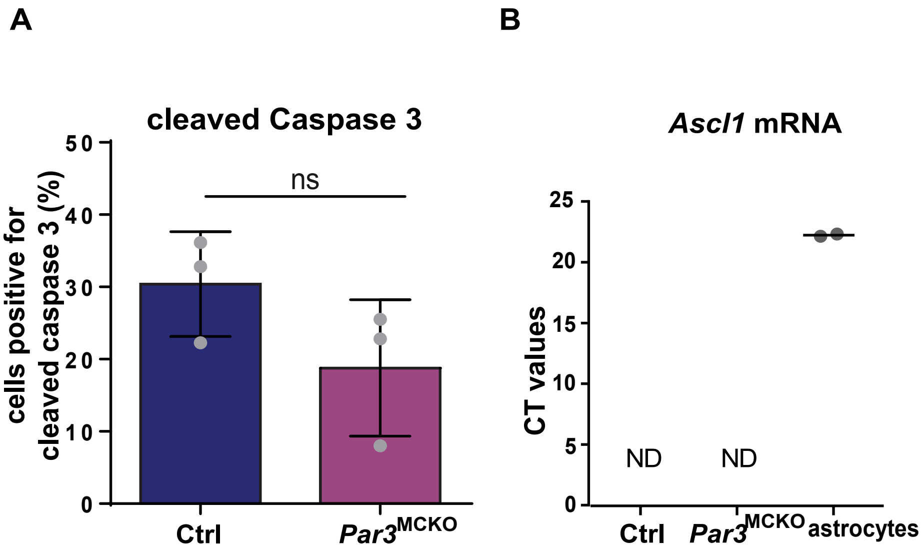
A
cleaved Caspase 3
cells positive for cleaved caspase 3 (%)
ns
0
10
20
30
40
50
Ctrl
Par3MCKO
B
Ascl1 mRNA
CT values
0
5
10
15
20
25
ND
ND
Ctrl
Par3MCKO
astrocytes

**Figure S3. No signs of increased apoptosis and altered lineage following Par3 deletion in melanocytes. Related to Figure 3. A)** Quantification of percentage of cleaved caspase 3-positive primary melanocytes isolated from P2 *Par3*$^{MCKO}$ and control mice immunostained for cleaved caspase 3, n=3, unpaired student t-test, mean±SD, ns: p=0.1658 **B)** CT values of RT-qPCR of *Ascl1* expression of primary melanocytes isolated from P2 *Par3*$^{MCKO}$ and control mice, n=3, ND=not detected. Astrocytes served as positive control.

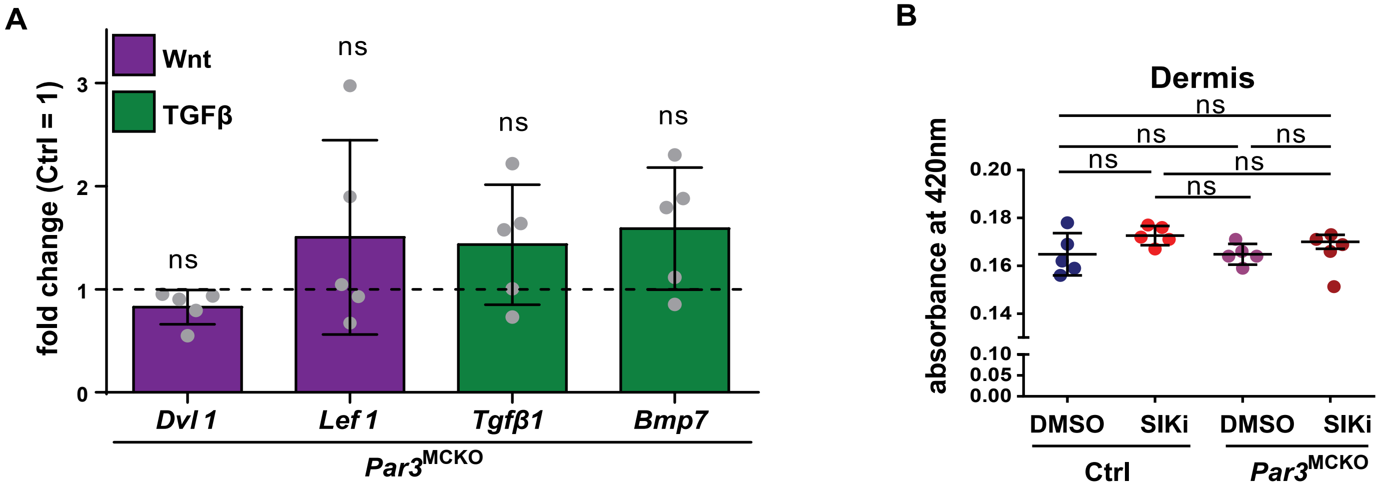
A
Wnt
TGFβ
fold change (Ctrl = 1)
0
1
2
3
ns
ns
ns
ns
Dvl 1
Lef 1
Tgfβ1
Bmp7
Par3MCKO
B
Dermis
absorbance at 420nm
0.20
0.18
0.16
0.14
0.10
0.05
0.00
ns
ns
ns
ns
ns
ns
DMSO
SIKi
DMSO
SIKi
Ctrl
Par3MCKO

**Figure S4. Expression of Wnt and TGFβ signaling targets is not altered in *Par3*$^{KO}$ melanocytes. Related to Figure 4. A)** Quantification of RT-qPCR of *Dvl1, Lef1, Tgfβ1, Bmp7* expression of primary melanocytes isolated from P2 *Par3*$^{MCKO}$ and control mice, n=5, one-sample t-test, mean±SD, *Dvl1*: ns: p>0.1, *Lef1*: ns: p>0.1 *Tgfβ1*: ns: p>0.1, *Bmp7*: ns: p>0.1. **B)** Melanin content assay. Quantification of spectrophotometry at 420nm of anterior tail skin dermis treated with DMSO or SIKi from *Par3*$^{MCKO}$ and control mice, n=5, one-way ANOVA, mean±SD, ns: p=0.1545 (Ctrl DMSO vs Ctrl SIKi), ns: p=0.999 (Ctrl DMSO vs *Par3*$^{MCKO}$ DMSO), ns: p=0.4638 (Ctrl DMSO vs *Par3*$^{MCKO}$ SIKi), ns: p=0.1545 (Ctrl SIKi vs *Par3*$^{MCKO}$ DMSO), ns: p=0.8767 (Ctrl SIKi vs *Par3*$^{MCKO}$ SIKi), ns: p=0.4638 (*Par3*$^{MCKO}$ DMSO vs *Par3*$^{MCKO}$ SIKi). Abbreviations: Ctrl, control; a.u., arbitrary units; P, postnatal day, SIKi, SIK inhibitor.

**Table S1. Antibodies and imaging reagents used in this study. Related to Experimental Procedures.**

| **Primary antibodies (supplier)** | **Catalog nr.** | **Clone/Ref.** | **Lot nr.** | **Species** | **Dilution** |
|---|---|---|---|---|---|
| cKit (Cell Signalling) | 3074 | D13A2 | 2 | rabbit | WB: 1:1000<br>IF: 1:100 |
| Cleaved caspase 3 (R&D Systems) | 9664 | 3/13 | 14 | rabbit | IF: 1:500 |
| GAPDH (Millipore) | MAB374 | 6C5 | 2322571 | mouse | WB 1:10.000 |
| MITF (Thermo scientific) | MA5-14146 | C5 | PL1945451 | mouse | IF:1:300 |
| Par3 (Millipore) | 07-330 | Polyclonal | 2615671 | rabbit | WB:<br>1:10.000 |
| Tyr (Santa Cruz) | sc-7833 | Polyclonal | N/A | goat | WB: 1:1000 |
| Trp2 (Santa Cruz) | sc-10451 | Polyclonal | N/A | goat | IF 1:300 |
| **Secondary antibodies for immunofluorescence analyses** | | | | | |
| AlexaFluor 488 α-mouse (Invitrogen) | A21202 | Polyclonal | 1741782 | donkey | IF 1:500 |
| AlexaFluor 488 α-rabbit (Invitrogen) | A21206 | Polyclonal | 1910751 | donkey | IF 1:500 |
| AlexaFluor 694 α-goat (Invitrogen) | A11058 | Polyclonal | N/A | donkey | IF 1:500 |
| **Secondary antibodies for immunoblot analyses** | | | | | |
| HRP α-rabbit (GE Healthcare) | NA9340V | Polyclonal | 9720820 | donkey | WB 1:4000 |
| HRP α-mouse (GE Healthcare) | NA931V | Polyclonal | 11076057 | sheep | WB 1:4000 |
| HRP α-goat (Invitrogen) | 81-1620 | Polyclonal | 1324727A | rabbit | WB 1:4000 |
| **Other Reagents** | | | | | |
| DAPI (Roth) | D1306 | N/A | N/A | N/A | 2μg/ml |

**Table S2. TaqMan® Gene Expression Assays (ThermoFisher Scientific) used in this study. Related to Experimental Procedures.**

| Target gene | ID | Cat. No. | label |
|---|---|---|---|
| *cKit* | Mm00445212_m1 | 4448892 | Fam |
| *Dvl1* | Mm00438592_m1 | 4448892 | Fam |
| *Gapdh* | Mm99999915_g1 | 4448490 | Vic |
| *Hprt* | Mm00446968_m1 | 4448489 | Vic |
| *Lef1* | Mm00550265_m1 | 4453320 | Fam |
| *Mitf* | Mm00434954_m1 | 4448892 | Fam |
| *Par3* | Mm00473929_m1 | 4448892 | Fam |
| *Tgfß1* | Mm01178820_m1 | 4453320 | Fam |
| *Tyr* | Mm00495817_m1 | 4448892 | Fam |
| *Tyrp1* | Mm00453201_m1 | 4448892 | Fam |

**Table S3. Primers used in this study. Related to Experimental Procedures.**

| **Name** | **Purpose** | **Sequence 5'-3'** |
|---|---|---|
| Cre-3 | Cre/ Genotyping | CGA TGC AAC GAG TGA TGA GGT TC |
| Cre-5 | Cre/ Genotyping | GCA CGT TCA CCG GCA TCA AC |
| +598iR | Par3A deletion PCR/ Genotyping | TAC CGT TAA CTG CAG CTC GGC TCT G |
| 0509iR | Par3A deletion PCR/ Genotyping | AGC TGG CGC TGG TAC CAT CTC CTC C |
| -423iF | Par3A deletion, flox PCR/ Genotyping | AGG CTA GCC TGG GTG ATT TGA GAC C |
| -159iR | Par3A flox PCR/ Genotyping | TTC CCT GAG GCC TGA CAC TCC AGT C |
